# Early-Emerging and Highly-Heritable Sensitivity to Human Communication in Dogs

**DOI:** 10.1101/2021.03.17.434752

**Authors:** Emily E. Bray, Gitanjali E. Gnanadesikan, Daniel J. Horschler, Kerinne M. Levy, Brenda S. Kennedy, Thomas R. Famula, Evan L. MacLean

## Abstract

Dogs exhibit similarities to humans in their sensitivity to cooperative-communicative cues, but the extent to which they are biologically prepared for communication with humans is heavily debated. To investigate the developmental and genetic origins of these traits, we tested 375 eight-week-old dog puppies on a battery of social-cognitive measures. We hypothesized that if dogs’ social skills for cooperating with humans are biologically prepared, then these skills should emerge robustly in early development, not require extensive socialization or learning, and exhibit heritable variation. Puppies were highly skillful at using diverse human gestures and we found no evidence of learning across test trials, suggesting that they possess these skills prior to their first exposure to these cues. Critically, over 40% of the variation in dogs’ point-following abilities and attention to human faces was attributable to genetic factors. Our results suggest that these social skills in dogs emerge early in development and are under strong genetic control.

**Highlights:**

- Genetic factors account for nearly half of variation in dog social skills
- Puppies displayed social skills and interest in human faces from 8 weeks old
- Puppies successfully used human gestures from the very first trial

---

Human cognition is believed to be unique in part due to early-emerging social skills that enable flexible forms of cooperative communication (*1*). Comparative studies show that at 2.5 years of age, human children reason about the physical world similarly to other great apes, yet already possess cognitive skills for cooperative communication far exceeding those found in our closest primate relatives (*2, 3*). Despite these differences between humans and other apes, a growing body of research indicates that domestic dogs are similar to human children in their sensitivity to cooperative-communicative acts. From early in development, dogs flexibly respond to diverse forms of cooperative gestures (*4, 5*). Like human children, dogs are sensitive to ostensive signals marking gestures as communicative, as well as contextual factors required for inferences about these communicative acts (*6–8*). Studies comparing dogs and wolves suggest that these traits may be evolutionarily derived in dogs, possibly as a response to similar selective pressures in human evolution and dog domestication (*6, 9, 10*). However, key questions about potential biological bases for these abilities remain untested.

For a trait to evolve under selection, the trait must a) vary between individuals, b) convey fitness benefits, and c) have a heritable basis (*11*). Regarding the first criterion, dogs, like human children, exhibit individual differences in correlated skills for cooperative communication, providing phenotypic variation upon which selection could act (*12*). Evidence for the second criterion of differential fitness is impossible to assess directly given the historical nature of dog domestication. However, studies with experimentally domesticated canids reveal changes in social cognition that arise as a byproduct of selection for tamability, a phenotype believed to have been targeted during dog domestication (*13*). Lastly, a heritable basis for social cognition in dogs has been suggested indirectly through comparative studies with wolves (*14, 15*) and across breeds (*16*), but direct evidence for genetic contributions to these traits remain elusive. Therefore, there is a critical need for research that can directly assess genetic contributions to these aspects of dog social cognition and inform their potential for evolutionary selection.

We hypothesized that if dogs’ social skills for cooperating with humans are biologically prepared (*17*), they should emerge robustly in early development, not require extensive socialization or learning, and exhibit heritable variation. To evaluate these predictions, we tested 375 retriever puppies from a pedigreed population (mean age = 8.5 weeks) on a cognitive and behavioral test battery. This battery was designed to broadly characterize the cognitive abilities of young puppies prior to extensive socialization with humans (*18*). With respect to social cognition, we included measures that assessed spontaneous responses to gestural communication, social approach and interaction with humans, and attention to human faces.

Before the gesture-following tasks, puppies completed a series of warm-up trials ensuring that they were motivated to search for a hidden food reward. In gesture-following test trials, a handler held the puppy in a central position, equidistant from two hiding locations. The experimenter provided ostensive cues by saying “puppy, look!” while initiating eye contact, and either pointed to and gazed at the location containing the reward (Fig 1A; pointing), or showed the puppy a small yellow block and placed it next to the baited container (Fig 1B; communicative marker). The experimenter then remained motionless while the puppy was released to search. To confirm that puppies could not locate the hidden reward using olfaction, we also conducted a series of control trials with an identical procedure, except the experimenter did not provide a social cue prior to the dog’s search. In the human-interest task, the experimenter stood outside the testing arena, looked at the puppy, and recited a standardized script using dog-directed speech [mimicking the prosody of infant-directed speech] (*19*). While speaking, the experimenter timed the duration of the puppy’s gaze to their face. The experimenter then entered the testing arena and petted the puppy if they approached within arm’s distance. The time that the puppy spent in proximity to the experimenter was recorded (*20*). Lastly, in the unsolvable task, puppies initially learned to displace the lid from a container to obtain a food reward within; then, on test trials, the lid to the container was fixed in place, rendering the problem unsolvable, and the experimenter timed the duration of the puppy’s gaze to their face (*21*).

**Fig 1.**
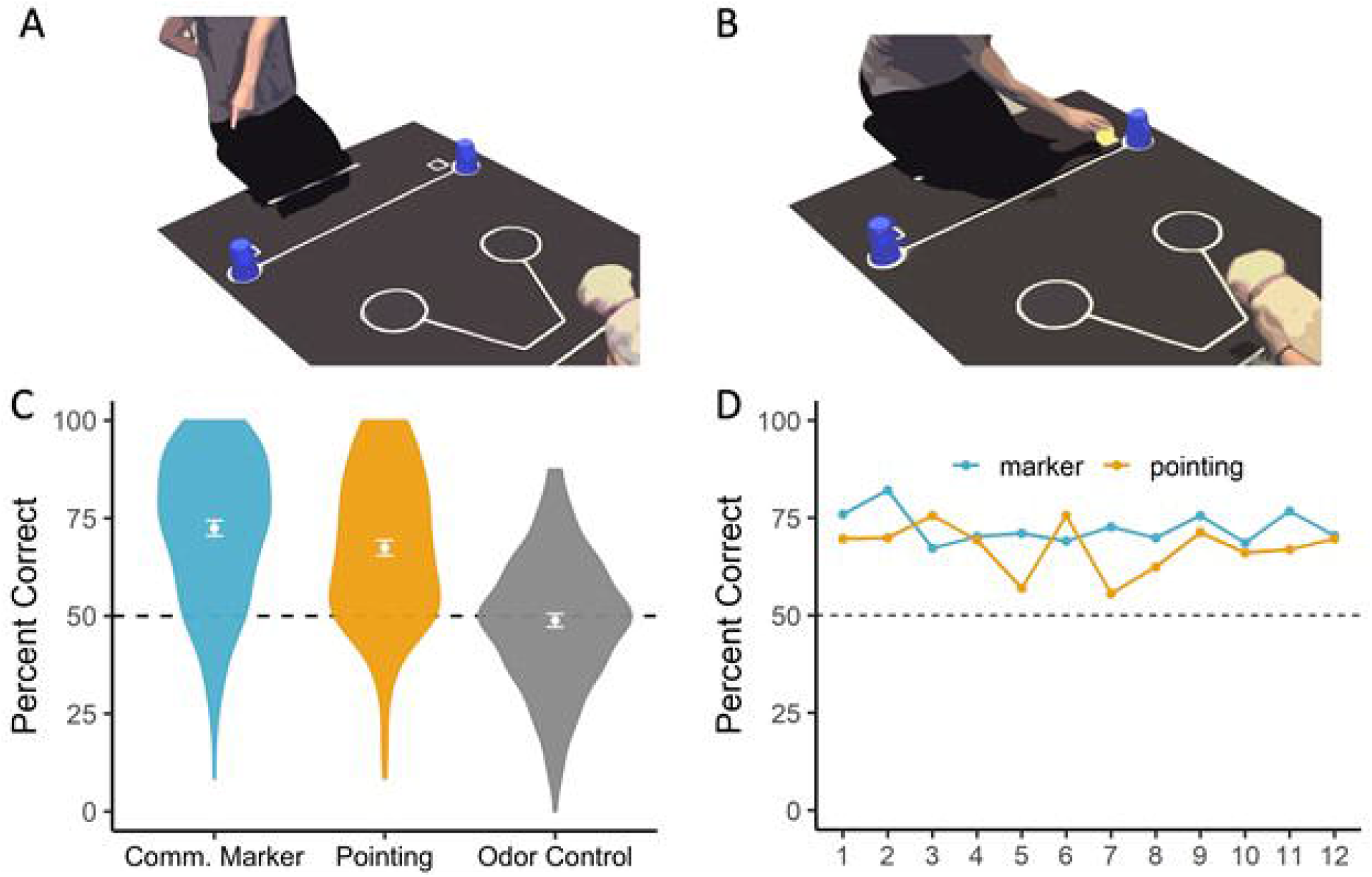
Procedure and results from the gesture-following tasks. The experimenter hid food in one of two locations and either, (A) pointed to and gazed at, or (B) placed an arbitrary marker next to, the baited container. (C) Puppies exceeded chance expectation with both social cues, but not in an olfactory control condition. Points and error bars reflect the mean and 95% confidence intervals. (D) Performance in the pointing and communicative marker tasks was above chance expectation from the first test trial, with no evidence for learning across trials.

In the gesture-following tasks, puppies reliably followed communicative cues at levels far exceeding chance expectation (Fig 1C; Pointing: mean = 67.4%, 95% CI = 65.5-69.3%; t_364_ = 17.58, p < 0.001, Cohen’s d = 0.92; Marker: mean = 72.4%, 95% CI = 70.5-74.38%; t_368_ = 22.88, p < 0.001, Cohen’s d = 1.19). However, in odor control trials, performance was not above chance expectation (mean = 48.9%, 95% CI = 47.2-50.5%; t_361_ = −1.33, p = 0.19, Cohen’s d = 0.07), confirming that success in the gesture-following tasks was not due to unintentional olfactory cueing. Importantly, in the gesture-following tasks, puppies performed above chance expectation from the first test trial (Pointing: accuracy = 70%, p < 0.001; Marker: accuracy 76%, p < 0.001; binomial tests), and showed no improvement in performance across trials (Fig 1D; Pointing: β_trial_ = −0.003, t_4014_ = −1.57, p = 0.12; Marker: β_trial_ = −0.003, t_4058_ = −1.36, p = 0.18). Therefore, from early in development and prior to extensive socialization with humans, dogs exhibit a robust sensitivity to human gestural communication that does not rely on learning. Puppies also spent time gazing at the experimenter as they were spoken to using dog-directed speech (mean = 6.2s gaze at face, 95% CI = 5.8-6.7s out of a possible 30s) and approached and interacted with this person when given the opportunity (mean = 18s interaction, 95% CI = 17.7-19.4s). On the other hand, although puppies looked to the human during the unsolvable task, they exhibited much less social gaze in this context (mean = 1.1s, 95% CI = 0.9-1.2s). These findings suggest that, like human children (*22, 23*), puppies excel at comprehending and responding to human-initiated social signals, while production of communicative behavior occurs later in development (*20*).

To assess a potential genetic basis for variation in these traits, we estimated their narrow-sense heritability using Bayesian linear mixed models incorporating a relatedness matrix for the study population (*24*), while controlling for breed, sex, age, and rearing location (Fig 2). As a statistical measure, heritability refers to the proportion of variance in a trait that is attributable to additive genetic factors. In humans, a meta-analysis of fifty years of twin studies suggests that on average, 47% of inter-individual variation on cognitive measures is attributable to genetic factors (*25*). However, much less is known about the heritability of cognitive traits in nonhuman animals, despite widespread agreement about its importance for understanding cognitive evolution (*26–28*). Among the five social measures, sensitivity to pointing gestures and attention to human faces during speech were estimated to have the highest heritability. In the point-following task, genetic factors accounted for 43% of variation between dogs (90% *h*^*2*^ credible interval = 0.20-0.68). Social attention toward the experimenter’s face while speaking to the subject had an equally high heritability estimate (mean *h*^*2*^ = 0.43; 90% *h*^*2*^ credible interval = 0.17-0.70), although the tendency to approach and interact with the experimenter in this context was less heritable (mean *h*^*2*^ = 0.13; 90% *h*^*2*^ credible interval = 0.00-0.34). Sensitivity to the communicative marker cue was less heritable than point following, but genetic factors still accounted for 14% of variation between puppies (mean *h*^*2*^ = 0.14; 90% *h*^*2*^ credible interval = 0.01-0.32). Lastly, the tendency to gaze at a human’s face during an unsolvable task was estimated to be the least heritable trait (mean *h*^*2*^ = 0.08, 90% *h*^*2*^ credible interval = 0.00-0.26). Notably however, compared to adult dogs, puppies exhibited very little social looking in this context (*20*), resulting in minimal phenotypic variance to be explained.

**Fig 2.**
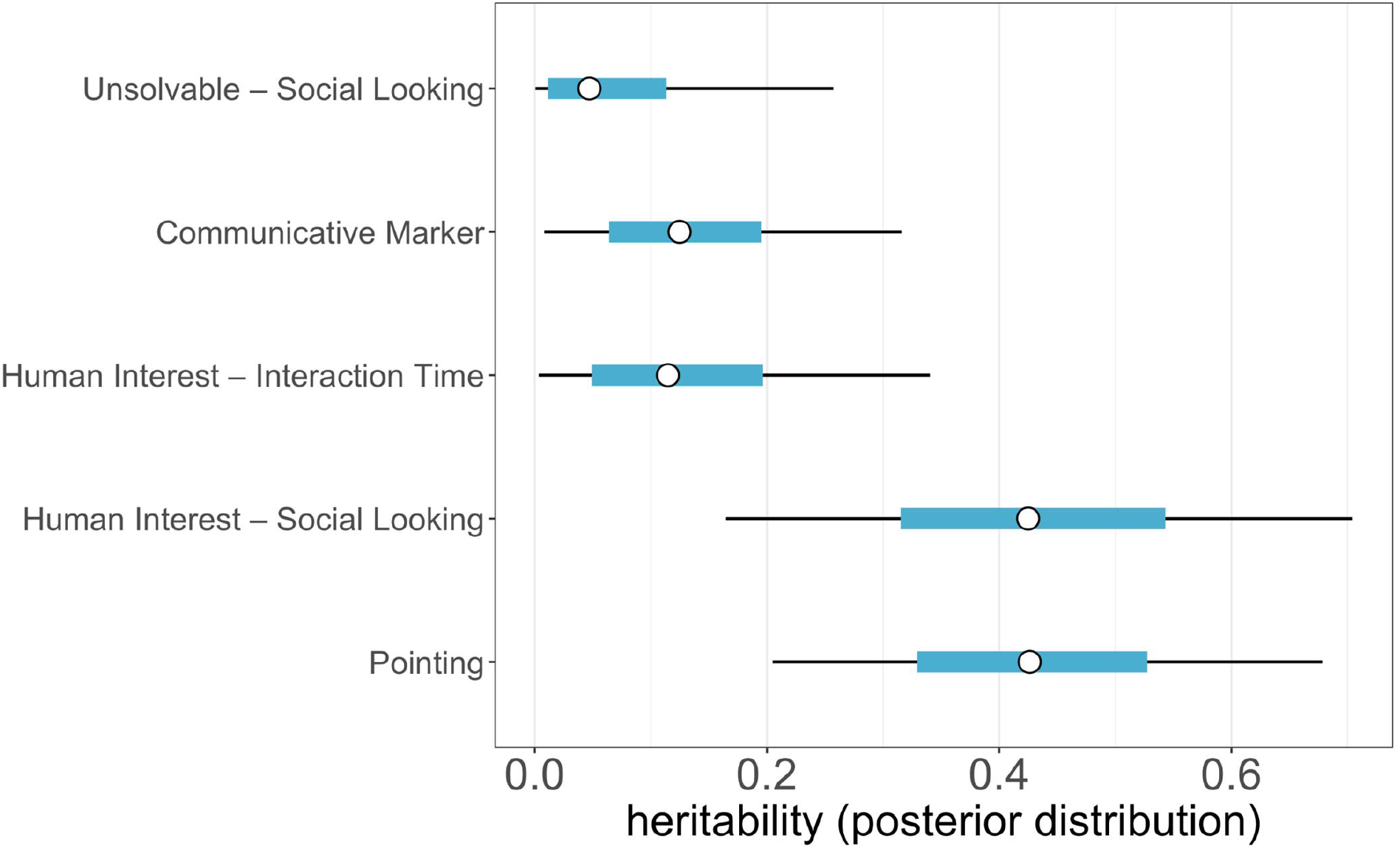
Posterior distributions of heritability estimates. Circles reflect the mean of the posterior distribution and thick blue and thin black bars span the 50% and 90% credible intervals, respectively.

Our findings show that from early in development dog puppies are highly sensitive and receptive to diverse communicative signals from humans, including gestures and speech, and that variation in these traits is under strong genetic control. Our study design also controls for several alternative explanations. First, subjects were tested at ~8 weeks, when they were still living with their littermates and eating, sleeping, and spending most of their time with conspecifics rather than humans. Despite their limited experience with humans, puppies were highly skilled at following human gestures and motivated to attend to and interact with humans. Second, our sample size of 375 puppies permitted a powerful analysis of potential learning effects during gesture-following tasks. These analyses confirmed that puppies were skillful from the very first test trial, and that their performance did not improve across trials. Third, puppies performed no better than chance expectation in odor control trials, confirming they were unable to locate the hidden rewards using olfaction.

A limitation of this work is that it does not identify the specific cognitive mechanism(s) supporting puppies’ use of human communicative cues. Although previous studies with dog puppies (*29*) and adults (*6, 30, 31*) suggest that dogs are sensitive to the ostensive nature of these signals and their communicative context, and that performance does not depend on “lower level” processes such as local enhancement, the mechanisms enabling dogs to interpret human communication remain debated (*32–37*). Importantly however, the significance of our findings is not tied to questions about mechanism. Rather, from a functional perspective, our study reveals that from early in development, individual dogs vary in their response to human communicative cues, and these differences are under strong genetic control. Regardless of the cognitive mechanisms involved, these heritable individual differences shape dogs’ responses to human communication and have potential to undergo selection. Similarly, our results do not suggest that a genetic basis for these traits is unique to dogs. Given that diverse species exhibit some sensitivity to human gestural communication (e.g., *38, 39*), it will be important for future work to assess potential genetic contributions to these traits in other lineages.

Collectively, our results demonstrate that dog social skills emerge robustly in early development and that variation in these traits is strongly influenced by genetic factors. Until now, researchers have relied on an untested assumption that variation in dog social cognition was heritable, rendering it unclear to what extent these traits might be available to selection. Our findings provide the first direct evidence that a large fraction of variation in dog social cognition is indeed heritable, and that this variation is expressed during a developmental window when historical selection pressures may have been particularly strong (*40, 41*). Although heritability estimates are specific to a given population, environment, and point in time, if similar heritable variation was present in the wolf populations that gave rise to dogs, these phenotypes would have had strong potential to undergo rapid selection. Given that genetic similarity among breeds also predicts variation in similar social-communicative processes (*16*), we hypothesize that genetic variation contributing to these phenotypes has relatively ancient origins and continues to contribute to variance in socio-cognitive skills, both within and among modern dog breeds. Our findings also set the stage for future research on the genetic architecture of these traits, an approach that has already proven fruitful for identifying common genetic underpinnings for other aspects of social behavior in dogs and humans (*42, 43*). Considering our results, we expect that dogs will provide a similarly powerful model for questions about the biological bases of the cooperative-communicative skills which are foundational to human social cognition.

## Acknowledgements

We thank Ben Allen, Erika Albrecht, Ashtyn Bernard, Molly Byrne, Elizabeth Carranza, Averill Cantwell, Mary Chiang, Amanda Chira, Alexzia Clark, Laura Douglas, Kyla Guinon, Erin Hardin, Theresa Hatcher, Victoria Holden, Emily Humphrey, Julia Kemper, Lindsey Lang, Camden Olson, Facundo Ortega, Gianna Ossello, Alessandra Ostheimer, Amber Robello, Camila Risueno-Pena, Ashley Ryan, Holland Smith, Paige Smith and Mia Wesselkamper for help with data collection and coding. We are also grateful to the staff of Canine Companions for Independence and their dedicated volunteer breeder caretakers for accommodating research with their assistance dog puppies at the Canine Early Development Center. We thank Brian Hare and Margaret Gruen for their help developing and piloting the measures used in this study.

## Author contributions

All authors contributed to Writing – Review & Editing. EB: Conceptualization, Methodology, Investigation, Data Curation, Writing – Original Draft, Supervision, Project administration, Funding acquisition. GG: Conceptualization, Methodology, Investigation, Data Curation, Project administration. DH: Conceptualization, Methodology, Investigation. KL: Methodology, Data Curation, Resources, Project administration. BK: Methodology, Resources, Supervision. TF: Software, Formal analysis. EM: Conceptualization, Methodology, Formal analysis, Writing – Original Draft, Visualization, Supervision, Funding acquisition.

## Materials and Methods

### Statistical Analyses

All statistical analyses were carried out in R v.3.6.0 (*44*). Behavioral variables were scored live, but all tasks were video recorded for reliability assessment. Independent coders scored from video all trials for ~20% of randomly selected subjects, and interrater reliability was calculated using Pearson correlation for continuous variables and Cohen’s kappa for categorial variables. There was high inter-rater agreement on all measures (Human Interest – Social Looking: *r* = 0.92; Human Interest – Interaction Time: *r* = 0.99; Unsolvable – Social Looking: *r* = 0.89; Communicative marker: κ = 0.97; Arm pointing: κ = 0.99; Odor control: κ = 0.97).

We estimated heritability using a Bayesian implementation of the animal model. The animal model is a linear mixed model that incorporates a random genetic effect with a known covariance structure for relatedness between individuals. Phenotypic variance (variation on the cognitive measures) is partitioned to determine the proportion of variance attributable to additive genetic effects. Relatedness between subjects was calculated from a pedigree for the study population.

Statistical models were fit using rstan (*45*) with weakly informative priors and included terms for breed (Labrador Retriever, Golden Retriever, or Lab-Golden cross), sex, age, and rearing location (volunteer Breeder Caretaker home or Canine Early Development Center). We applied rank-based inverse normal transformation for all dependent measures. Models were implemented using four separate chains, each of which was run for 75,000 iterations using the no-U-turn sampler (NUTS) algorithm. The first 5,000 iterations were used as a burn-in, after which we employed a thinning interval of 20 draws between samples retained for the posterior distribution. The final posterior distributions consisted of samples merged across the four independent chains. Rhat values indicated that chains converged successfully in all models (all Rhat values < 1.01). The effective sample sizes and 90% credible intervals for the heritability estimates in all models are reported in Table S1.

### Methods

#### Subjects

All subjects belonged to Canine Companions for Independence (CCI), the largest nonprofit provider of assistance dogs for people with disabilities in the United States. Previous research has shown that, as adults, this population performs comparably to a heterogenous population of pet dogs on similar cooperative-communicative measures (*11*). Specifically, when participating in an arm pointing and communicative marker task, adult performance level did not significantly differ between the two populations (pointing: t_156.64_ = −1.25, *p* = 0.21; marker: t_118.19_ = 1.72, *p* = 0.09). On the pointing task, pet dogs (N = 87) chose correctly on 67.74% of trials and assistance dogs (N = 194) chose correctly on 64.18% of trials. On the marker task, pet dogs (N = 79) chose correctly on 80.80% of trials and assistance dogs (N = 189) chose correctly on 85.45% of trials. CCI provided informed consent to all aspects of the study, and all testing procedures were reviewed and adhered to regulations set forth by the Institutional Animal Care and Use Committee at the University of Arizona (IACUC No. 16-175).

We tested 375 puppies (203 females and 172 males) from February 2017 through June 2020 when they were approximately 8 weeks of age (range 7.3-10.4 weeks; mean = 8.4 weeks), prior to being placed with volunteer puppy raisers. Our sample consisted of 254 Labrador x golden crosses, 98 Labrador retrievers and 23 golden retrievers from 117 different litters (Table S2). While all puppies successfully completed at least one task and the vast majority of puppies successfully completed all four of the social-cognitive tasks plus the odor control condition, there were a few subjects who were unable to complete every task (see below for refamiliarization/abort criteria). Table S3 indicates the final sample size for each task.

Puppies were either whelped and weaned in the homes of local volunteer breeder caretakers (N = 261) or at the Canine Early Development Center (CEDC; N = 114), a professional facility with full-time technicians who monitor and care for the dams and puppies. After birth, puppies remained with their mother until weaning was complete, around six weeks of age, and then were housed with their littermates until being sent to live with their respective puppy raisers. During this time, puppies spend most of their time eating, sleeping, and playing with conspecifics. At any given moment the dog-human ratio is quite high, whereas once puppies are sent to their puppy-raising family the dog-human ratio is much lower, with each dog then receiving much more individualized attention and training. Regardless of birth environment, all puppies spent time at the CEDC between 7.5-10 weeks of age to receive a veterinary examination and routine vaccinations prior to their puppy raising placement. All puppies participated in cognitive testing at the CEDC before being placed with their volunteer puppy raiser.

#### Procedures

All procedures followed previously published protocols from our earlier studies. Testing took place over a series of three days in sessions that lasted approximately 45 minutes – 1 hour at a time for each puppy. Puppies participated in hiding-finding warm-ups on days 1 through 3, the human interest task on day 1, and the unsolvable, communicative marker, arm pointing, and odor control tasks on day 2. For a comprehensive overview of this test battery (the Dog Cognitive Development Battery), we direct the reader to Bray et al. (*17, 19*). Descriptions of general testing procedures, and the methods for the social-cognitive tasks presented in this paper, are detailed below. The below text is adapted and reprinted from Animal Behaviour, 166, Bray EE, Gruen ME, Gnanadesikan GE, Horschler DJ, Levy KM, Kennedy BS, Hare BA, MacLean EL, Cognitive characteristics of 8- to 10-week-old assistance dog puppies, 193-206, Copyright (2020) with permission from Elsevier.

#### General Methods

Puppies were tested in a 5.9 × 4.3 m room at the CEDC of CCI, where the testing arena was enclosed by a 1.8 × 3 m free-standing exercise pen. Brown noise was played in the background during all testing to mask the sound of stopwatches and any ambient noise. For the communicative marker and arm pointing tasks, kibble was taped to the inside bottom of both containers as a control for odor cues. The mat was marked with starting lines for the experimenter and the subject. Testing took place between 0800 and 2030 hours. During testing, puppies were either housed together at the CEDC (N = 339) or housed at the breeder caretaker’s home who then transported them to the CEDC each morning (N = 36). We withheld the majority of each subject’s meal preceding the session, and withheld kibble was then soaked and used as the reward during testing. The experimenter and the dog handler were always in the room with the puppy except during the ‘human interest’ task, when the handler left the room to minimize distractions. All puppies were naïve to all tasks. Testing sessions were videotaped using Canon video cameras (Vixa HF R700) with wide-angle lenses (Pearstone 43 mm, Vivitar HD MC AF High Definition) mounted on tripods.

#### Handler Guidelines

The handler’s interactions with the puppy were limited to positioning the puppy at the start line and releasing the puppy at the beginning of each trial. At the beginning of each trial, the handler (hereafter H) sat or kneeled at the starting position, centered the subject in the starting box and looked straight down (Fig. S1). The puppy was always held by the shoulders, with two hands placed evenly on either side, or by the back of the collar. Once the experimenter (hereafter E) gave the ‘Okay!’ release signal, H released the puppy. If the subject did not immediately leave the starting box, E would continue to repeat the release command as H reoriented or lifted the puppy by the haunches to encourage forward movement. As soon as the subject left the starting box, H looked up.

#### Choice criteria for object-choice tasks

Object-choice tasks included hiding–finding warm-ups, communicative marker, arm pointing, and odor control. In these tasks, a choice was defined as the subject physically touching the cup with their snout or a front paw. For each trial, the puppy was allowed up to 25 s from their release to make a choice. When a choice was made, H said ‘Choice’. If E saw the choice first, despite looking down, E could also say ‘Choice’. If the subject chose the baited cup, E praised the subject and lifted the cup so the puppy could eat the food reward before H repositioned the subject in the starting box. If the subject chose the incorrect cup, E said ‘Wrong’ in a neutral, monotone voice and the puppy was not given a food reward. E then returned the subject to H and retrieved the reward from the correct location before the next trial was administered. If the puppy did not make a choice within 25 s, the trial was repeated until the puppy made a choice or until the maximum number of repeated trials had been conducted (see below: Refamiliarization/Abort criteria).

#### Refamiliarization/Abort criteria

For hiding–finding warm-ups, if there were eight no choices (NCs) total in either phase 2 (one-cup alternating visible displacement) or phase 3 (two-cup visible displacement), including refamiliarizations, the task was aborted. When 12 trials of phase 2 or 20 trials of phase 3 were conducted (including repeated trials, but not refamiliarizations) and the pass criterion had not been met, the task was aborted. If the subject completed 20 trials of phase 3 but made the correct choice on at least the last two trials, more trials were conducted until the puppy made an incorrect choice or the pass criterion was met.

For the unsolvable task, if the puppy did not touch the container on two consecutive trials, the food reward was changed. If the subject did not finish all four trials within 12 attempts at familiarization trials, the task was aborted.

For object-choice tasks (i.e., communicative marker, arm pointing, odor control), if the subject did not make a choice within 25 s, the trial was repeated. If the puppy did not choose twice in a row, two trials of two-cup alternating hiding–finding warm-ups were performed. If at any point there were two more consecutive NCs, the food reward was increased and the puppy was refamiliarized with two more trials of two-cup hiding–finding. If the puppy did not engage in refamiliarization trials or made eight NCs in a single task, the task was aborted.

If at any point the subject became sleepy or uninterested in participating, a break was taken during which the puppy was given a chance to eliminate and E and H attempted to re-engage the puppy with play. In addition to not meeting task-specific demands, a task was aborted and the session was stopped if the puppy showed any signs of not feeling well (e.g., refusing to eat kibble, diarrhea, vomiting).

Aborted tasks were attempted again later in the same day, with a break of at least 30 min. If these tasks were aborted again, they were attempted for a final time in an extra session at the end of the battery, time permitting.

#### Hiding-finding warm-ups

Warm-up trials were meant to ensure that the subjects were motivated to search for the reward and to prevent side biases. These trials consisted of three phases: (1) no-cup visible placement and free-form cup familiarization game, (2) one-cup alternating visible placement, and (3) two-cup visible placement. The subject was required to pass a warm-up criterion prior to completing any other object-choice task.

##### Phase 1 – no-cup visible placement and free-form cup game

E presented the reward, then visibly placed the food on the mat in two locations: first halfway between the puppy and E, and then directly in front of E. When E placed the food on the mat, she said ‘Puppy, look!’. E then rested with her hands behind her back, looked straight down and said ‘Okay!’. The subject was allowed to approach the food and obtain the reward. If the subject did not approach the kibble within 25 s, the trial was repeated. If the puppy approached/climbed on E or sat and waited for E, the trial was repeated until the puppy retrieved the kibble immediately on their own.

After the puppy retrieved the visible food successfully from each location, E played a free-form cup game with the puppy to familiarize them to finding food under cups. The puppy watched as E placed food under a single cup, then was rewarded when they touched the cup. As needed, E tilted or tapped the cup to draw the puppy’s attention to the food and encourage the puppy to touch the cup. These hiding instances happened in quick succession to keep the puppy fully engaged. This activity was continued until puppies were reliably approaching and touching the cup.

##### Phase 2 – one-cup alternating

E presented the reward and visibly baited a single cup (lifting the cup straight up, ~10 cm off the ground) placed in either the right or left position (the position on the opposite side remained empty). After baiting, E kneeled at the starting location in the resting position. The subject was then allowed to approach the cup and touch it to obtain the reward. If the subject did not approach the cup within 25 s, E called the puppy over, showed the kibble if necessary, and rewarded them for touching the cup; the trial was then repeated. This phase of warm-ups familiarized the subject with the set-up and ensured that the subject was motivated to find the reward. Repeating NC trials served as a correction procedure for spontaneous side biases and ensured that subjects gained experience finding the reward in both locations. The subject was required to successfully retrieve the reward on four trials, twice on each side, to move on (within a maximum of 12 trials).

##### Phase 3 – two-cup visible displacement

Two cups were placed in the small circles connected to the 1 m line (Fig. S1). E presented the reward to the puppy and visibly baited one of the two cups. The baited location was predetermined and counterbalanced between locations in blocks of 10 trials, and the same side was never baited on more than two trials in a row. After baiting, E kneeled in the resting position. The subject was then allowed 25 s to make a choice. If the subject chose the baited cup, the subject was allowed to have the reward, E praised the subject, and the next trial was administered. If the subject chose the incorrect cup, E said ‘Wrong’ in a neutral tone, the subject was not rewarded, and the trial was repeated until the subject chose correctly. If the subject did not choose any cup within 25 s, the trial was repeated. This phase of warm-ups ensured that the subject was not choosing cups randomly and was attending to the experimenter’s actions. Subjects were required to choose correctly on their first attempt in four out of five consecutive trials, within a maximum of 20 attempts, to advance to test trials (with the caveat that if the subject reached 20 trials but made the correct choice on at least the last two trials, more trials were conducted until the puppy made an incorrect choice or the pass criterion was met).

#### Human interest

E stood inside the pen until at least 10 s after H left the room. E then exited the pen and stood 20 cm from the corner of the pen nearest the door. E’s left foot was perpendicular to the corner of the pen, and E’s right foot was facing the edge of the pen.

##### Phase 1 – test trials

E looked directly at the puppy, made eye contact whenever the puppy attended to her and recited the following script using dog-directed speech, which took approximately 30 s. E used the silent count-up timer to measure the amount of eye contact made during this time:

‘Hi pup! Are you a good puppy? Yes you are. What a good puppy. Aww, look how cute you are. Look at those big eyes and floppy ears. You’re such a cute puppy! Do you like to play? Are these experiments fun? You’re coming back tomorrow to play with me! We’ll play more cup games. We’re gonna have so much fun. I can’t wait to play with you! Are you the best puppy? Yes, you are. Of course you are. That’s a good puppy!’

##### Phase 2 – 30 s play break

E then started a 30 s count-down timer, stepped into the center of the pen and faced the direction she came from. If the puppy approached E during this time, E said ‘Hi pup!’, bent down, and petted the puppy as long as they were within reach, and was then quiet for the rest of the time. E did not move until the timer elapsed.

E repeated phases 1 and 2 in succession three times. The dependent measures for this task were the number of seconds that the puppy made eye contact with the human during phase 1 averaged across the three trials and the number of seconds that the puppy interacted with the human during phase 2 averaged across the three trials. For a subset of the sample (N = 15), puppies only completed two trials of phase 2 and thus, for these subjects, the number of seconds spent interacting with the human during phase 2 was only averaged over two trials. The overall mean was the same when removing these puppies from analysis.

#### Unsolvable task

The puppy was held by H at a distance of 100 cm from the center of a transparent container (Rubbermaid Brilliance small container; 10 × 14 cm and 6 cm high) affixed to a board (60 × 60 cm and 1.5 cm high). E was centered and behind the apparatus, directly behind the board, to allow room for the puppy to walk around the container. The reward for this task was three pieces of soaked kibble per trial.

##### Familiarization trials

In familiarization trials, E presented the subject with the reward, placed the reward inside the clear container and positioned the lid upside down and loosely on top of the container. Four warm-up trials were conducted with the following conditions:

1. The lid was propped on the side, not covering the container.
2. The lid covered ~½ of the container.
3. The lid covered ~ ¾ of the container.
4. The lid (again) covered ~ ¾ of the container.

After baiting the apparatus, E looked at the container and said ‘Okay!’ H released the puppy and started a 30 s timer. The puppy was allowed to approach the container, displace the lid and obtain the reward. The trial ended when the puppy retrieved the kibble. If the puppy did not retrieve the reward at the end of the 30 s trial, E called them over, drew their attention to the container and ensured that the puppy ate the kibble. That trial was then repeated. The puppy was required to successfully complete four familiarization trials without help before advancing to the test.

##### Test trials

Test trials were identical to familiarization trials except that E snapped the lid onto the container so that it could not be removed, thus rendering the problem unsolvable, and looked at the puppy as soon as she gave the release command. Subjects were given 30 s to attempt to access the reward. E remained seated and visually followed the puppy (rotating her torso if necessary) during this period. E held a silent stopwatch behind her back and started and stopped it in count-up mode to measure the total time that the puppy looked to her face. H timed the 30 s trial and looked down throughout the trial, ignoring the puppy. After each test trial, E praised the puppy, drew their attention to the container, opened the container and allowed the puppy to retrieve the contents. Four test trials were conducted. The dependent measure for this task was the number of seconds that the puppy made eye contact with the human averaged over all four test trials.

#### Gesture following (Communicative Marker and Arm Pointing)

E placed two cups centered on the line connecting the final cup locations and then placed the occluder (0.5 x 60 cm and 18 cm high) about 10 cm in front of the cups, blocking the puppy’s view of the cups and subsequent baiting. E presented the subject with the reward, placed the treat behind one of the cups (but near the center), lifted both cups and placed them over the treat and empty space. E then removed the occluder, placing it outside of the pen and slid the cups into the positions marked on the mat while looking downward.

##### Communicative marker

The marker (yellow wooden block) was initially placed in front of the cups while baiting. After baiting as described above, E picked up the marker in her right hand, leaned forward to show it to the subject (~10 cm from the subject’s nose) and said ‘Puppy, look!’. E then pulled the marker back to the central position, lifted it up and made eye contact with the puppy and said ‘Puppy, look!’ before placing the marker next to the appropriate cup (in the square marked on the mat; Fig. S1). After placing the marker, E looked down at a central point between the cups and held this position until a choice was made. E then gave an ‘Okay!’ release, signaling H to release the subject to make a choice. Until this point, H was holding the puppy but looking down so that H was unaware which side was being indicated. Once E gave the ‘Okay!’ release signal and the puppy left the starting box, H looked up. E maintained the resting position until the subject made a choice, or the maximum trial time of 25 s had elapsed. Twelve trials were conducted, and the dependent measure for this task was the percentage of trials that a puppy chose the cup next to which E placed the wooden block.

##### Arm pointing

After baiting as above, E made eye contact with the subject, said ‘Puppy, look!’ and pointed towards the cup with the reward. The pointing gesture consisted of the index finger of the contralateral hand extended about 20 cm from the cup with E’s head and gaze directed towards the cup. E then gave an ‘Okay!’ release, signaling H to release the subject to make a choice. Until this point, H was holding the puppy but looking down so that H was unaware which side was being indicated. Once E had given the ‘Okay!’ release signal and the puppy had left the starting box, H looked up; when the puppy made a choice, H said ‘Choice’. E maintained the static pointing gesture and gazed towards the correct cup until the subject made a choice or the maximum trial time of 25 s had elapsed. Twelve trials were conducted, and the dependent measure was the percentage of trials that a subject chose the cup towards which E gestured.

##### Odor control

Non-baited, clean cups were used for this task, and no cue was administered during the eight odor control trials. After baiting as above, E kneeled in the resting position and maintained her gaze down at a central point between the cups, holding this position until a choice was made. E then said ‘Okay!’, signaling H to release the subject to make a choice. Eight trials were conducted and the dependent measure for this task was the percentage of trials that a puppy chose the baited cup.

## Supplemental information

Tables S1 - S3

