## Supplemental Table 1 for "Early-Emerging and Highly-Heritable Sensitivity to Human Communication in Dogs"

Table S1.

Descriptive statistics for the posterior distributions of heritability estimates.

| ***trait*** | ***mean*** | ***sd*** | ***5%*** | ***50%*** | ***95%*** | ***effective sample size*** |
| --- | --- | --- | --- | --- | --- | --- |
| Pointing | 0.43 | 0.14 | 0.20 | 0.43 | 0.68 | 8,472 |
| Communicative Marker | 0.14 | 0.10 | 0.01 | 0.12 | 0.32 | 12,399 |
| Human Interest – Social Looking | 0.43 | 0.16 | 0.17 | 0.42 | 0.70 | 7,434 |
| Human Interest – Interaction Time | 0.13 | 0.11 | 0.00 | 0.12 | 0.34 | 11,258 |
| Unsolvable – Social Looking | 0.08 | 0.09 | 0.00 | 0.05 | 0.26 | 10,756 |
