## Supplemental Table 2 for "Early-Emerging and Highly-Heritable Sensitivity to Human Communication in Dogs"

Table S2.

Subject demographics.

| ***subject*** | ***sex*** | ***breed*** | ***litter*** | ***age at first day of testing (in weeks)*** |
| --- | --- | --- | --- | --- |
| 1 | Female | Labrador–golden retriever cross | 1 | 10.14 |
| 2 | Male | Labrador–golden retriever cross | 1 | 10.14 |
| 3 | Female | Labrador–golden retriever cross | 1 | 10.14 |
| 4 | Male | Labrador–golden retriever cross | 1 | 10.14 |
| 5 | Female | Labrador–golden retriever cross | 2 | 10.00 |
| 6 | Male | Labrador–golden retriever cross | 2 | 10.00 |
| 7 | Female | Labrador retriever | 3 | 9.29 |
| 8 | Female | Labrador retriever | 3 | 9.29 |
| 9 | Female | Labrador retriever | 3 | 9.29 |
| 10 | Female | Labrador–golden retriever cross | 4 | 9.14 |
| 11 | Male | Labrador–golden retriever cross | 4 | 9.14 |
| 12 | Male | Labrador–golden retriever cross | 4 | 9.14 |
| 13 | Female | Labrador retriever | 5 | 9.43 |
| 14 | Female | Labrador retriever | 5 | 9.43 |
| 15 | Male | Labrador retriever | 5 | 9.43 |
| 16 | Female | Labrador–golden retriever cross | 6 | 9.43 |
| 17 | Female | Labrador–golden retriever cross | 6 | 9.43 |
| 18 | Male | Labrador–golden retriever cross | 6 | 9.43 |
| 19 | Male | Labrador–golden retriever cross | 7 | 10.00 |
| 20 | Female | Labrador–golden retriever cross | 7 | 10.00 |
| 21 | Male | Labrador retriever | 8 | 9.43 |
| 22 | Female | Labrador retriever | 8 | 9.43 |
| 23 | Female | Labrador–golden retriever cross | 9 | 9.71 |
| 24 | Male | Labrador–golden retriever cross | 9 | 9.71 |
| 25 | Female | Labrador–golden retriever cross | 10 | 9.86 |
| 26 | Male | Labrador–golden retriever cross | 10 | 9.86 |
| 27 | Female | Labrador–golden retriever cross | 11 | 9.86 |
| 28 | Female | Labrador–golden retriever cross | 11 | 9.86 |
| 29 | Female | Labrador retriever | 12 | 9.71 |
| 30 | Female | Labrador retriever | 12 | 9.71 |
| 31 | Female | Labrador–golden retriever cross | 13 | 9.86 |
| 32 | Male | Labrador–golden retriever cross | 13 | 9.86 |
| 33 | Male | Labrador–golden retriever cross | 13 | 9.86 |
| 34 | Female | Labrador–golden retriever cross | 14 | 10.00 |
| 35 | Male | Labrador–golden retriever cross | 14 | 10.00 |
| 36 | Male | Labrador–golden retriever cross | 15 | 9.57 |
| 37 | Female | Labrador–golden retriever cross | 15 | 9.57 |
| 38 | Male | Labrador–golden retriever cross | 16 | 9.43 |
| 39 | Male | Golden retriever | 16 | 9.43 |
| 40 | Female | Labrador–golden retriever cross | 16 | 9.43 |
| 41 | Female | Labrador–golden retriever cross | 16 | 9.43 |
| 42 | Female | Labrador–golden retriever cross | 17 | 10.00 |
| 43 | Female | Labrador–golden retriever cross | 17 | 10.00 |
| 44 | Male | Labrador–golden retriever cross | 18 | 9.29 |
| 45 | Female | Labrador–golden retriever cross | 18 | 9.29 |
| 46 | Female | Labrador–golden retriever cross | 18 | 9.29 |
| 47 | Female | Labrador retriever | 19 | 9.14 |
| 48 | Male | Labrador retriever | 19 | 9.14 |
| 49 | Female | Labrador–golden retriever cross | 20 | 9.43 |
| 50 | Female | Labrador–golden retriever cross | 20 | 9.43 |
| 51 | Male | Labrador–golden retriever cross | 21 | 8.57 |
| 52 | Female | Labrador–golden retriever cross | 21 | 8.57 |
| 53 | Male | Labrador retriever | 22 | 8.86 |
| 54 | Female | Labrador retriever | 22 | 8.86 |
| 55 | Female | Labrador–golden retriever cross | 23 | 9.14 |
| 56 | Male | Labrador–golden retriever cross | 23 | 9.14 |
| 57 | Male | Labrador–golden retriever cross | 24 | 10.43 |
| 58 | Female | Labrador–golden retriever cross | 24 | 10.43 |
| 59 | Female | Labrador–golden retriever cross | 24 | 10.43 |
| 60 | Male | Labrador–golden retriever cross | 25 | 9.43 |
| 61 | Female | Labrador–golden retriever cross | 25 | 9.43 |
| 62 | Female | Labrador–golden retriever cross | 25 | 9.43 |
| 63 | Female | Labrador retriever | 26 | 9.57 |
| 64 | Male | Labrador retriever | 26 | 9.57 |
| 65 | Female | Labrador–golden retriever cross | 27 | 9.14 |
| 66 | Female | Golden retriever | 28 | 9.00 |
| 67 | Male | Golden retriever | 28 | 9.00 |
| 68 | Female | Labrador–golden retriever cross | 28 | 9.00 |
| 69 | Male | Labrador–golden retriever cross | 29 | 9.29 |
| 70 | Female | Labrador–golden retriever cross | 29 | 9.29 |
| 71 | Female | Labrador retriever | 30 | 9.00 |
| 72 | Male | Labrador retriever | 30 | 9.00 |
| 73 | Female | Labrador–golden retriever cross | 31 | 8.57 |
| 74 | Male | Labrador–golden retriever cross | 31 | 8.57 |
| 75 | Male | Labrador retriever | 32 | 9.71 |
| 76 | Female | Labrador retriever | 32 | 9.71 |
| 77 | Female | Labrador retriever | 32 | 9.71 |
| 78 | Male | Labrador–golden retriever cross | 33 | 9.29 |
| 79 | Male | Labrador–golden retriever cross | 33 | 9.29 |
| 80 | Female | Labrador–golden retriever cross | 33 | 9.29 |
| 81 | Male | Labrador–golden retriever cross | 33 | 9.71 |
| 82 | Female | Labrador–golden retriever cross | 33 | 9.71 |
| 83 | Female | Labrador retriever | 34 | 9.29 |
| 84 | Female | Labrador–golden retriever cross | 35 | 9.14 |
| 85 | Female | Labrador–golden retriever cross | 35 | 9.14 |
| 86 | Male | Labrador–golden retriever cross | 35 | 9.14 |
| 87 | Female | Golden retriever | 36 | 9.29 |
| 88 | Female | Golden retriever | 36 | 9.29 |
| 89 | Male | Labrador–golden retriever cross | 37 | 9.14 |
| 90 | Female | Golden retriever | 37 | 9.14 |
| 91 | Male | Labrador–golden retriever cross | 38 | 9.00 |
| 92 | Female | Labrador–golden retriever cross | 38 | 9.14 |
| 93 | Female | Labrador retriever | 39 | 9.43 |
| 94 | Male | Labrador retriever | 39 | 9.43 |
| 95 | Male | Labrador–golden retriever cross | 40 | 9.43 |
| 96 | Female | Labrador–golden retriever cross | 40 | 9.43 |
| 97 | Female | Labrador–golden retriever cross | 41 | 9.29 |
| 98 | Male | Labrador–golden retriever cross | 41 | 9.29 |
| 99 | Male | Labrador–golden retriever cross | 41 | 9.29 |
| 100 | Female | Labrador–golden retriever cross | 42 | 9.14 |
| 101 | Male | Labrador–golden retriever cross | 42 | 9.14 |
| 102 | Female | Labrador–golden retriever cross | 43 | 9.71 |
| 103 | Male | Labrador–golden retriever cross | 43 | 9.71 |
| 104 | Male | Labrador–golden retriever cross | 44 | 9.71 |
| 105 | Male | Labrador–golden retriever cross | 44 | 9.71 |
| 106 | Female | Labrador–golden retriever cross | 44 | 9.71 |
| 107 | Female | Labrador retriever | 45 | 9.00 |
| 108 | Female | Labrador retriever | 46 | 9.00 |
| 109 | Male | Labrador retriever | 46 | 9.00 |
| 110 | Male | Labrador retriever | 46 | 9.00 |
| 111 | Female | Labrador–golden retriever cross | 47 | 9.14 |
| 112 | Female | Labrador–golden retriever cross | 47 | 9.14 |
| 113 | Female | Labrador–golden retriever cross | 48 | 9.14 |
| 114 | Male | Labrador–golden retriever cross | 48 | 9.14 |
| 115 | Male | Labrador–golden retriever cross | 49 | 8.71 |
| 116 | Female | Labrador–golden retriever cross | 49 | 8.71 |
| 117 | Female | Labrador–golden retriever cross | 50 | 8.71 |
| 118 | Male | Labrador–golden retriever cross | 50 | 8.71 |
| 119 | Female | Labrador–golden retriever cross | 50 | 8.71 |
| 120 | Female | Labrador–golden retriever cross | 51 | 9.00 |
| 121 | Male | Labrador–golden retriever cross | 51 | 9.00 |
| 122 | Female | Labrador–golden retriever cross | 51 | 9.00 |
| 123 | Male | Labrador retriever | 52 | 8.71 |
| 124 | Male | Labrador retriever | 52 | 8.71 |
| 125 | Female | Labrador retriever | 52 | 8.71 |
| 126 | Male | Labrador–golden retriever cross | 53 | 8.86 |
| 127 | Female | Labrador–golden retriever cross | 53 | 8.86 |
| 128 | Male | Labrador–golden retriever cross | 53 | 8.86 |
| 129 | Female | Labrador–golden retriever cross | 53 | 8.86 |
| 130 | Female | Labrador–golden retriever cross | 54 | 8.71 |
| 131 | Male | Labrador–golden retriever cross | 54 | 8.71 |
| 132 | Female | Labrador–golden retriever cross | 54 | 8.71 |
| 133 | Male | Labrador retriever | 55 | 9.29 |
| 134 | Male | Labrador retriever | 55 | 9.29 |
| 135 | Female | Labrador retriever | 55 | 9.29 |
| 136 | Female | Labrador retriever | 55 | 9.29 |
| 137 | Female | Labrador retriever | 56 | 9.57 |
| 138 | Male | Labrador retriever | 56 | 9.57 |
| 139 | Male | Labrador retriever | 56 | 9.57 |
| 140 | Male | Labrador–golden retriever cross | 57 | 8.86 |
| 141 | Female | Labrador–golden retriever cross | 57 | 8.86 |
| 142 | Female | Labrador–golden retriever cross | 57 | 8.86 |
| 143 | Male | Labrador–golden retriever cross | 58 | 8.86 |
| 144 | Female | Labrador–golden retriever cross | 58 | 8.86 |
| 145 | Male | Labrador–golden retriever cross | 59 | 8.71 |
| 146 | Female | Labrador–golden retriever cross | 59 | 8.71 |
| 147 | Female | Labrador–golden retriever cross | 59 | 8.71 |
| 148 | Female | Labrador–golden retriever cross | 59 | 8.71 |
| 149 | Male | Labrador–golden retriever cross | 60 | 8.57 |
| 150 | Female | Labrador–golden retriever cross | 60 | 8.57 |
| 151 | Female | Labrador–golden retriever cross | 60 | 8.57 |
| 152 | Male | Labrador–golden retriever cross | 61 | 8.57 |
| 153 | Female | Labrador–golden retriever cross | 61 | 8.57 |
| 154 | Female | Labrador–golden retriever cross | 61 | 8.57 |
| 155 | Male | Labrador retriever | 62 | 8.86 |
| 156 | Female | Labrador retriever | 62 | 8.86 |
| 157 | Male | Labrador–golden retriever cross | 63 | 8.43 |
| 158 | Male | Labrador–golden retriever cross | 63 | 8.43 |
| 159 | Female | Labrador–golden retriever cross | 63 | 8.43 |
| 160 | Female | Labrador–golden retriever cross | 63 | 8.43 |
| 161 | Female | Labrador–golden retriever cross | 64 | 8.00 |
| 162 | Male | Labrador–golden retriever cross | 64 | 8.00 |
| 163 | Female | Labrador–golden retriever cross | 64 | 8.00 |
| 164 | Male | Labrador–golden retriever cross | 64 | 8.00 |
| 165 | Female | Labrador–golden retriever cross | 65 | 7.86 |
| 166 | Female | Labrador–golden retriever cross | 65 | 7.86 |
| 167 | Male | Labrador–golden retriever cross | 65 | 7.86 |
| 168 | Male | Labrador–golden retriever cross | 65 | 7.86 |
| 169 | Male | Labrador-golden retriever cross | 66 | 7.71 |
| 170 | Female | Labrador-golden retriever cross | 66 | 7.71 |
| 171 | Female | Labrador-golden retriever cross | 66 | 7.71 |
| 172 | Female | Labrador-golden retriever cross | 66 | 7.71 |
| 173 | Female | Labrador-golden retriever cross | 67 | 8.43 |
| 174 | Female | Labrador-golden retriever cross | 67 | 8.43 |
| 175 | Male | Labrador-golden retriever cross | 67 | 8.43 |
| 176 | Female | Labrador-golden retriever cross | 67 | 8.43 |
| 177 | Male | Labrador-golden retriever cross | 67 | 8.43 |
| 178 | Male | Labrador-golden retriever cross | 68 | 8.00 |
| 179 | Male | Labrador-golden retriever cross | 68 | 8.00 |
| 180 | Female | Labrador-golden retriever cross | 68 | 8.00 |
| 181 | Female | Labrador-golden retriever cross | 68 | 8.00 |
| 182 | Male | Labrador retriever | 69 | 7.43 |
| 183 | Female | Labrador retriever | 69 | 7.43 |
| 184 | Female | Labrador retriever | 69 | 7.43 |
| 185 | Female | Labrador retriever | 69 | 7.43 |
| 186 | Female | Labrador retriever | 69 | 7.43 |
| 187 | Male | Labrador retriever | 70 | 8.14 |
| 188 | Female | Labrador retriever | 70 | 8.14 |
| 189 | Male | Labrador retriever | 70 | 8.14 |
| 190 | Female | Labrador retriever | 70 | 8.14 |
| 191 | Female | Labrador-golden retriever cross | 71 | 8.57 |
| 192 | Male | Labrador-golden retriever cross | 71 | 8.57 |
| 193 | Male | Labrador-golden retriever cross | 71 | 8.57 |
| 194 | Male | Labrador-golden retriever cross | 71 | 8.57 |
| 195 | Female | Golden retriever | 72 | 8.14 |
| 196 | Female | Golden retriever | 72 | 8.14 |
| 197 | Male | Golden retriever | 72 | 8.14 |
| 198 | Male | Golden retriever | 72 | 8.14 |
| 199 | Male | Labrador-golden retriever cross | 73 | 8.57 |
| 200 | Male | Labrador-golden retriever cross | 73 | 8.57 |
| 201 | Female | Labrador-golden retriever cross | 73 | 8.57 |
| 202 | Female | Labrador-golden retriever cross | 73 | 8.57 |
| 203 | Female | Labrador retriever | 74 | 7.71 |
| 204 | Male | Labrador retriever | 74 | 7.71 |
| 205 | Female | Labrador retriever | 74 | 7.71 |
| 206 | Male | Labrador retriever | 74 | 7.71 |
| 207 | Female | Labrador-golden retriever cross | 75 | 8.00 |
| 208 | Male | Labrador-golden retriever cross | 75 | 8.00 |
| 209 | Female | Labrador-golden retriever cross | 75 | 8.00 |
| 210 | Male | Labrador-golden retriever cross | 75 | 8.00 |
| 211 | Male | Labrador-golden retriever cross | 76 | 8.57 |
| 212 | Female | Labrador-golden retriever cross | 76 | 8.57 |
| 213 | Male | Labrador-golden retriever cross | 76 | 8.57 |
| 214 | Female | Labrador-golden retriever cross | 76 | 8.57 |
| 215 | Male | Labrador retriever | 77 | 7.71 |
| 216 | Male | Labrador retriever | 77 | 7.71 |
| 217 | Female | Labrador retriever | 77 | 7.71 |
| 218 | Female | Labrador retriever | 77 | 7.71 |
| 219 | Male | Labrador-golden retriever cross | 78 | 8.00 |
| 220 | Female | Labrador-golden retriever cross | 78 | 8.00 |
| 221 | Female | Labrador-golden retriever cross | 78 | 8.00 |
| 222 | Female | Labrador-golden retriever cross | 78 | 8.00 |
| 223 | Male | Labrador-golden retriever cross | 79 | 8.14 |
| 224 | Male | Labrador-golden retriever cross | 79 | 8.14 |
| 225 | Male | Labrador-golden retriever cross | 79 | 8.14 |
| 226 | Female | Labrador-golden retriever cross | 79 | 8.14 |
| 227 | Female | Golden retriever | 80 | 7.86 |
| 228 | Male | Golden retriever | 80 | 7.86 |
| 229 | Female | Golden retriever | 80 | 7.86 |
| 230 | Male | Golden retriever | 80 | 7.86 |
| 231 | Female | Labrador-golden retriever cross | 81 | 8.14 |
| 232 | Female | Labrador-golden retriever cross | 81 | 8.14 |
| 233 | Male | Labrador-golden retriever cross | 81 | 8.14 |
| 234 | Female | Labrador-golden retriever cross | 81 | 8.14 |
| 235 | Male | Labrador-golden retriever cross | 82 | 7.86 |
| 236 | Male | Labrador-golden retriever cross | 82 | 7.86 |
| 237 | Female | Labrador-golden retriever cross | 82 | 7.86 |
| 238 | Female | Labrador-golden retriever cross | 82 | 7.86 |
| 239 | Female | Labrador retriever | 83 | 8.14 |
| 240 | Female | Labrador retriever | 83 | 8.14 |
| 241 | Male | Labrador retriever | 83 | 8.14 |
| 242 | Male | Labrador retriever | 83 | 8.14 |
| 243 | Male | Labrador retriever | 83 | 8.14 |
| 244 | Female | Labrador-golden retriever cross | 84 | 8.14 |
| 245 | Male | Labrador-golden retriever cross | 84 | 8.14 |
| 246 | Male | Labrador-golden retriever cross | 84 | 8.14 |
| 247 | Male | Labrador-golden retriever cross | 84 | 8.14 |
| 248 | Male | Labrador retriever | 85 | 7.86 |
| 249 | Female | Labrador retriever | 85 | 7.86 |
| 250 | Female | Labrador retriever | 85 | 7.86 |
| 251 | Female | Labrador retriever | 85 | 7.86 |
| 252 | Female | Labrador retriever | 86 | 7.86 |
| 253 | Male | Labrador retriever | 86 | 7.86 |
| 254 | Female | Labrador retriever | 86 | 7.86 |
| 255 | Male | Labrador retriever | 86 | 7.86 |
| 256 | Male | Labrador-golden retriever cross | 87 | 7.71 |
| 257 | Male | Labrador-golden retriever cross | 87 | 7.71 |
| 258 | Female | Labrador-golden retriever cross | 87 | 7.71 |
| 259 | Female | Labrador-golden retriever cross | 87 | 7.71 |
| 260 | Female | Labrador-golden retriever cross | 88 | 7.86 |
| 261 | Female | Labrador-golden retriever cross | 88 | 7.86 |
| 262 | Male | Labrador-golden retriever cross | 88 | 7.86 |
| 263 | Male | Labrador-golden retriever cross | 88 | 7.86 |
| 264 | Male | Labrador-golden retriever cross | 89 | 7.86 |
| 265 | Male | Labrador-golden retriever cross | 89 | 7.86 |
| 266 | Male | Labrador-golden retriever cross | 89 | 7.86 |
| 267 | Female | Labrador-golden retriever cross | 90 | 7.43 |
| 268 | Male | Labrador-golden retriever cross | 90 | 7.43 |
| 269 | Female | Labrador-golden retriever cross | 90 | 7.43 |
| 270 | Male | Labrador-golden retriever cross | 90 | 7.43 |
| 271 | Female | Labrador-golden retriever cross | 91 | 7.86 |
| 272 | Male | Labrador-golden retriever cross | 91 | 7.86 |
| 273 | Male | Labrador-golden retriever cross | 91 | 7.86 |
| 274 | Female | Labrador-golden retriever cross | 91 | 7.86 |
| 275 | Female | Labrador-golden retriever cross | 92 | 7.29 |
| 276 | Male | Labrador-golden retriever cross | 92 | 7.29 |
| 277 | Female | Labrador-golden retriever cross | 92 | 7.29 |
| 278 | Male | Labrador-golden retriever cross | 92 | 7.29 |
| 279 | Male | Labrador retriever | 93 | 8.57 |
| 280 | Female | Labrador retriever | 93 | 8.57 |
| 281 | Female | Labrador retriever | 93 | 8.57 |
| 282 | Male | Labrador retriever | 93 | 8.57 |
| 283 | Male | Golden retriever | 94 | 7.29 |
| 284 | Female | Golden retriever | 94 | 7.29 |
| 285 | Female | Golden retriever | 94 | 7.29 |
| 286 | Male | Golden retriever | 94 | 7.29 |
| 287 | Female | Golden retriever | 95 | 7.57 |
| 288 | Male | Golden retriever | 95 | 7.57 |
| 289 | Male | Golden retriever | 95 | 7.57 |
| 290 | Female | Labrador-golden retriever cross | 95 | 7.57 |
| 291 | Male | Labrador-golden retriever cross | 96 | 7.71 |
| 292 | Female | Labrador-golden retriever cross | 96 | 7.71 |
| 293 | Female | Labrador-golden retriever cross | 96 | 7.71 |
| 294 | Male | Labrador-golden retriever cross | 96 | 7.71 |
| 295 | Male | Labrador-golden retriever cross | 97 | 8.14 |
| 296 | Female | Labrador-golden retriever cross | 97 | 8.14 |
| 297 | Female | Labrador-golden retriever cross | 97 | 8.14 |
| 298 | Male | Labrador-golden retriever cross | 97 | 8.14 |
| 299 | Female | Golden retriever | 98 | 7.57 |
| 300 | Female | Golden retriever | 98 | 7.57 |
| 301 | Female | Labrador-golden retriever cross | 98 | 7.57 |
| 302 | Male | Labrador-golden retriever cross | 98 | 7.57 |
| 303 | Male | Labrador-golden retriever cross | 99 | 7.71 |
| 304 | Female | Labrador-golden retriever cross | 99 | 7.71 |
| 305 | Female | Labrador-golden retriever cross | 99 | 7.71 |
| 306 | Female | Labrador-golden retriever cross | 99 | 7.71 |
| 307 | Male | Labrador-golden retriever cross | 100 | 7.86 |
| 308 | Female | Labrador-golden retriever cross | 100 | 7.86 |
| 309 | Male | Labrador-golden retriever cross | 100 | 7.57 |
| 310 | Female | Labrador-golden retriever cross | 100 | 7.86 |
| 311 | Male | Labrador retriever | 101 | 7.57 |
| 312 | Female | Labrador retriever | 101 | 7.57 |
| 313 | Female | Labrador retriever | 101 | 7.57 |
| 314 | Male | Labrador retriever | 101 | 7.57 |
| 315 | Male | Labrador-golden retriever cross | 102 | 8.00 |
| 316 | Male | Labrador-golden retriever cross | 102 | 8.00 |
| 317 | Female | Labrador-golden retriever cross | 102 | 8.00 |
| 318 | Female | Labrador-golden retriever cross | 102 | 8.00 |
| 319 | Female | Labrador-golden retriever cross | 103 | 8.14 |
| 320 | Male | Labrador-golden retriever cross | 103 | 8.14 |
| 321 | Female | Labrador-golden retriever cross | 103 | 8.14 |
| 322 | Male | Labrador-golden retriever cross | 103 | 8.14 |
| 323 | Female | Labrador-golden retriever cross | 104 | 8.43 |
| 324 | Female | Labrador-golden retriever cross | 104 | 8.43 |
| 325 | Male | Labrador-golden retriever cross | 104 | 8.43 |
| 326 | Male | Labrador-golden retriever cross | 104 | 8.43 |
| 327 | Male | Labrador retriever | 105 | 7.86 |
| 328 | Male | Labrador retriever | 105 | 7.86 |
| 329 | Female | Labrador retriever | 105 | 7.86 |
| 330 | Male | Labrador retriever | 105 | 7.86 |
| 331 | Male | Labrador retriever | 106 | 7.71 |
| 332 | Male | Labrador retriever | 106 | 7.71 |
| 333 | Male | Labrador retriever | 106 | 7.71 |
| 334 | Female | Labrador retriever | 106 | 7.71 |
| 335 | Male | Labrador retriever | 107 | 8.71 |
| 336 | Female | Labrador retriever | 107 | 8.71 |
| 337 | Female | Labrador retriever | 107 | 8.71 |
| 338 | Male | Labrador retriever | 107 | 8.71 |
| 339 | Male | Labrador-golden retriever cross | 108 | 8.43 |
| 340 | Female | Labrador-golden retriever cross | 108 | 8.43 |
| 341 | Male | Labrador-golden retriever cross | 109 | 7.57 |
| 342 | Male | Labrador-golden retriever cross | 109 | 7.57 |
| 343 | Male | Labrador-golden retriever cross | 109 | 7.57 |
| 344 | Female | Labrador-golden retriever cross | 109 | 7.57 |
| 345 | Male | Labrador-golden retriever cross | 110 | 7.57 |
| 346 | Female | Labrador-golden retriever cross | 110 | 7.57 |
| 347 | Female | Labrador-golden retriever cross | 110 | 7.57 |
| 348 | Male | Labrador-golden retriever cross | 110 | 7.57 |
| 349 | Female | Labrador-golden retriever cross | 111 | 7.71 |
| 350 | Female | Labrador-golden retriever cross | 111 | 7.71 |
| 351 | Male | Labrador-golden retriever cross | 111 | 7.71 |
| 352 | Female | Labrador-golden retriever cross | 111 | 7.71 |
| 353 | Female | Labrador-golden retriever cross | 112 | 7.71 |
| 354 | Male | Labrador-golden retriever cross | 112 | 7.71 |
| 355 | Male | Labrador-golden retriever cross | 112 | 7.71 |
| 356 | Female | Labrador-golden retriever cross | 112 | 7.71 |
| 357 | Female | Labrador-golden retriever cross | 113 | 8.29 |
| 358 | Male | Labrador-golden retriever cross | 113 | 8.29 |
| 359 | Female | Labrador-golden retriever cross | 113 | 8.29 |
| 360 | Female | Labrador-golden retriever cross | 113 | 8.29 |
| 361 | Female | Labrador-golden retriever cross | 114 | 7.71 |
| 362 | Female | Labrador-golden retriever cross | 114 | 7.71 |
| 363 | Male | Labrador-golden retriever cross | 114 | 7.71 |
| 364 | Male | Labrador-golden retriever cross | 114 | 7.71 |
| 365 | Male | Labrador retriever | 115 | 7.43 |
| 366 | Female | Labrador retriever | 115 | 7.43 |
| 367 | Female | Labrador retriever | 115 | 7.43 |
| 368 | Male | Labrador retriever | 115 | 7.43 |
| 369 | Female | Labrador-golden retriever cross | 116 | 7.57 |
| 370 | Female | Labrador-golden retriever cross | 116 | 7.57 |
| 371 | Male | Labrador-golden retriever cross | 116 | 7.57 |
| 372 | Male | Labrador retriever | 117 | 7.71 |
| 373 | Male | Labrador retriever | 117 | 7.71 |
| 374 | Female | Labrador retriever | 117 | 7.71 |
| 375 | Female | Labrador retriever | 117 | 7.71 |
