## Supplemental Table 3 for "Early-Emerging and Highly-Heritable Sensitivity to Human Communication in Dogs"

Table S3.

Number of puppies that aborted each task, and the final sample size for each task.

| ***task*** | ***no. of puppies that aborted*** | ***final sample size*** |
| --- | --- | --- |
| Human Interest | 0 | 357 |
| Unsolvable | 0 | 375 |
| Communicative Marker | 6 | 369 |
| Arm Pointing | 10 | 365 |
| Odor Control | 13 | 362 |

Human interest was added shortly after data collection began but was successfully completed by all 357 puppies who were presented with it.
